# Evolution of seasonal plasticity in response to climate change differs between life-stages of a butterfly

**DOI:** 10.1101/2022.12.16.520735

**Authors:** Matthew E. Nielsen, Sören Nylin, Christer Wiklund, Karl Gotthard

## Abstract

Climate change alters seasonal environments without altering photoperiod, creating a cue-environment mismatch for organisms that rely on photoperiod as a cue for seasonal plasticity and phenology. Evolution can potentially correct for this mismatch by altering the photoperiodic reaction norm, but often phenology depends on multiple plastic decisions made at different life stages and times of year. We tested whether seasonal plasticity in different life stages evolves independently or in concert under climate change using *Pararge aegeria* (Speckled wood butterfly). This butterfly uses day length as a cue for life history plasticity in two different life stages: larval development time and pupal diapause. Photoperiodic reaction norms for plasticity in these traits were first measured over 30 years ago for two different Swedish populations. In this study, we replicated historic experiments that measured these reaction norms using the contemporary populations. We found evidence for evolution of the reaction norm for larval development time, but in opposite directions in the two populations. In contrast, we found no evidence for evolution of the reaction norm for pupal diapause. These results show that different life stages can evolve differently in response to climate change and only studying one part of the life cycle will not always be enough to fully understand how climate change impacts phenotypic plasticity and phenology.

## Introduction

Changes in phenology are one of the best documented examples of biological responses to climate change (1, 2); however, the actual magnitude of these changes and how they correspond to local climate change vary dramatically among taxa (2–4). When these shifts vary among interacting species, they can create phenological mismatches between those species, disrupting that interaction with severe ecological consequences (2–5). Despite their importance, the mechanisms underlying these shifts in phenology have rarely been documented, particularly whether they represent phenotypic plasticity, genetic evolution, or the evolution of plasticity (6–9).

One factor that may severely constrain the ability to adjust phenology in response to climate change is reliance on photoperiod as a cue for the life history plasticity that determines phenology. Particularly at higher latitudes, day length is commonly used as a reliable indicator of the time of year because it has historically been a strong predictor of the seasons themselves (10, 11). Nevertheless, photoperiod is one aspect of the environment that does not change with climate, and thus organisms that rely more heavily on photoperiod as a cue are predicted to adjust their phenology less under climate change than species that rely on other, climate-sensitive cues (12). This can lead to inappropriate plastic responses to the new climate, with potentially disastrous fitness consequences (13, 14). Evolution of the photoperiodic reaction norm for seasonal plasticity could correct for these mismatches and allow phenology to adjust to the new climate. Nevertheless, few studies have demonstrated contemporary evolution of photoperiodic plasticity in association with climate change (15–17).

Phenology, however, often depends on multiple plastic decisions made at different times of year and different stages in the life cycle, and this complicates the analysis of evolutionary and plastic responses to climate change (2). For example, the timing of migration in birds depends on molting, physiological development, and behavior—all of which occur in different seasons and can evolve separately (18). Do the reaction norms for seasonal plasticity in different life stages evolve in concert to synchronize the whole life cycle in relation to climate change, or do they instead evolve independently to account for different changes at different times of year?

*Pararge aegeria,* the speckled wood butterfly, is an excellent example of a species whose phenology depends on multiple plastic decisions across its life cycle, creating strong variation among populations in the annual number of generations (19, 20). Both larval and pupal development time are highly photoperiod sensitive and, depending on when during autumn they develop, they can enter winter diapause (delayed development) as either a pupa or third-instar larva (21). Further, photoperiodic plasticity in larval development rates is an important component of seasonal life cycle regulation in *P. aegeria:* they typically delay their larval development at intermediate photoperiods associated with late summer (22). This is proposed to ensure that they reach pupation at an appropriate time to enter pupal diapause rather than completing direct development to become an adult at a time when they cannot complete another generation (23). All of these decisions are primarily determined by photoperiod (23, 24), although modified slightly by temperature (20), meaning that *P. aegeria* should be strongly affected by the cue-environment mismatch created by rapid climate change. Nevertheless, there is extensive evidence for local adaptation of these reaction norms to natural spatial variation in photoperiod and climate (e.g., (20, 23)), which suggests that evolution might also be able to compensate for a climate change-induced mismatch—if it occurs quickly enough.

Here, we sought to test whether the photoperiodic reaction norms for the larval and pupal development time of *P. aegeria* have evolved over the last 30-35 years, as would be expected given climate change. We did so by taking advantage of a unique opportunity to perform an allochronic common garden experiment that compares life history plasticity in different life stages between the present and past. We replicated two historical experiments

(23, 24) that measured the photoperiodic reaction norms for these traits from 1985-1987 and in 1989, respectively. The original studies spanned an extremely broad range of photoperiods. Here we focused on a narrower subset of these photoperiods, including conditions that in the historical populations produced either direct development across the life cycle, delayed larval development associated with late summer, or pupal diapause. For both larval development time and pupal diapause time, we predicted a leftward shift of the reaction norm to allow the species to better take advantage of the extended growing seasons created by climate change (Figure 1). Further, these two studies also worked with populations from different parts of Sweden, with different seasonal life histories. The central Swedish population is univoltine while the southern Swedish population is bivoltine (20, 22, 25), but both have the capacity for the full range of developmental pathways (24). These are genetically distinct populations with a disjointed distribution (26, 27), allowing us to test whether evolutionary responses to climate change differ among populations from different parts of a species range with distinctly different phenologies and life histories.

**Figure 1.**
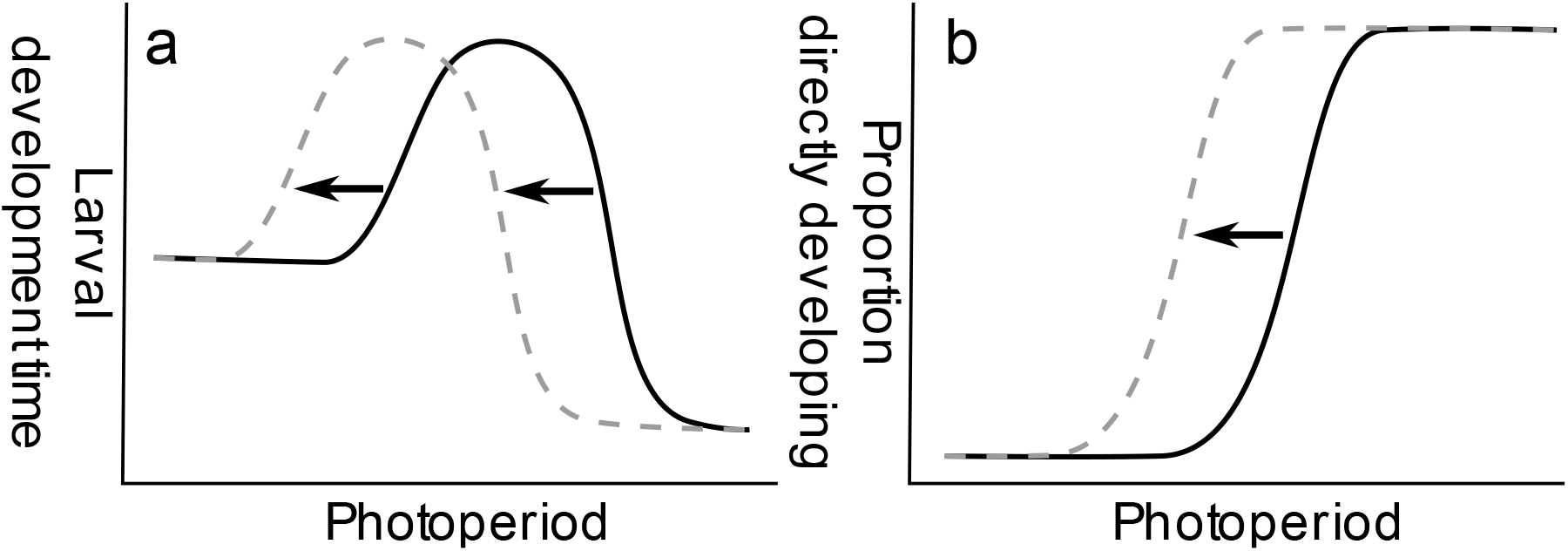
Hypotheses for evolutionary changes in the photoperiodic reaction norms for (a) larval development time and (b) pupal development pathway (diapause or direct development) associated with climate change. Warmer climates should lead to a longer growing season, and thus we predicted a leftward shift in both reaction norms to take advantage of that change. Solid black lines represent the approximate shape of the original reaction norm and dashed gray lines the predicted contemporary reaction norm following adaptation to climate change.

## Results

### Change in environmental temperatures between time periods

These two populations have experienced similar magnitudes of climate change, but the distribution of that change across the year differs between sites (Table S1, S2; Figure 2). Specifically, we compared seasonal temperature variation during the decade prior to each study using records from nearby weather stations. Mean annual temperatures increased by about 1.5°C at both field sites. Warming was generally greatest in winter and spring, but this seasonal pattern was much stronger for the central Swedish population. At the central site, warming was concentrated in winter (2.86°C), which warmed by 1.16°C more than the southern site (1.70°C). Across the rest of the year, warming was greater at the southern site, particularly summer during which the southern site warmed by 1.34°C, 0.47°C more than the central site (0.87°C). As a result, the estimated increase in growing season length was much greater for the southern population (31.4 days) than the central population (9.5 days). The start of the growing season advanced in both populations, but more in the south (southern: 20.4 days, central: 10.6 days), and the end of the growing season only changed meaningfully in in the south (southern: 11.0 days delay, central: 1.1 day advance).

**Figure 2.**
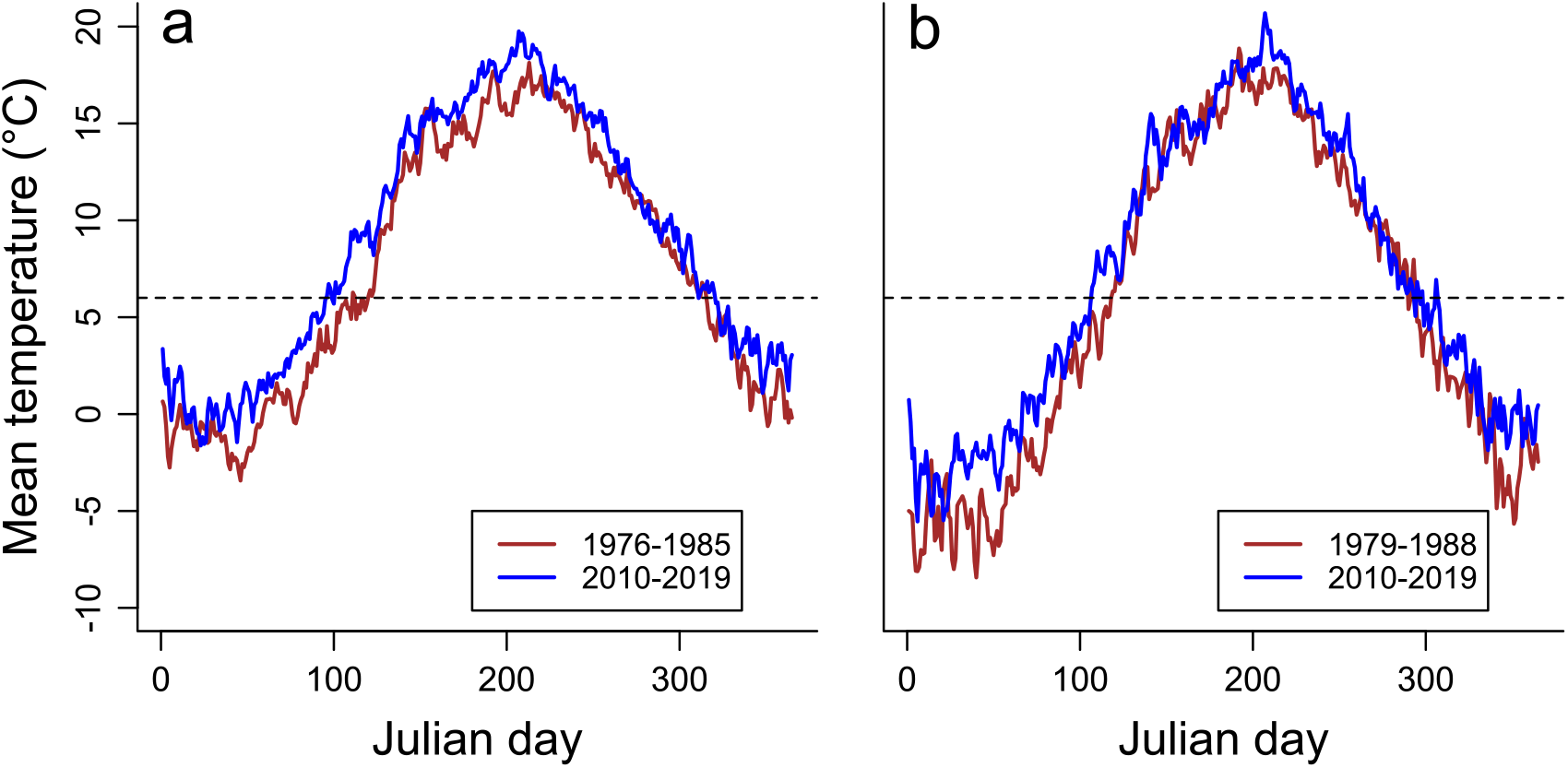
Change in mean daily temperatures during the decade before each study for the collection sites in (a) southern Sweden and (b) central Sweden. Historic data (red line) is from 1976-1985 for southern Sweden and 1979-1988 for central Sweden. Contemporary data (blue line) is from 2010-2019 for both sites. Dashed line represents 6°C, the minimum temperature for development in *P. aegeria* (20).

### Replication of historic experiments

To determine whether this climate change is associated with evolutionary changes in the seasonal plasticity of larval development time or pupal diapause, we repeated the historic experiments (23, 24) using the descendants of female *P. aegeria* collected in 2020 from the same locations as the original studies. For the southern population, we measured larval development time across four photoperiod treatments from 15L:9D to 18L:6D, comparing the results for 114 second-generation descendants of the contemporary population (2020) to 196 larvae measured in the historic experiments (1985-1987). We found a statistically clear interaction in which the response to photoperiod differed between the historic and contemporary studies (photoperiod x time=period: *F_3,294_*=7.72, *p*=0.000056, Figure 3A), but differently for different sexes (photoperiod x time-period x sex: *F_3,294_*=3.45, *p*=0.017). At most photoperiods, development time did not differ substantially between sexes or time period, and the main difference occurred at 17L:7D. Historically, this photoperiod represented a transition between direct and delayed development, with more males following this slower developmental pathway. However, development is faster in the contemporary study with nearly all individuals developing directly, and thus the difference between the sexes is likewise greatly reduced. In post-hoc tests, this is reflected in statistically clear differences between historic and contemporary males at 17L:7D and a clear difference between sexes at this photoperiod in the past, but not the present (Table S3).

**Figure 3.**
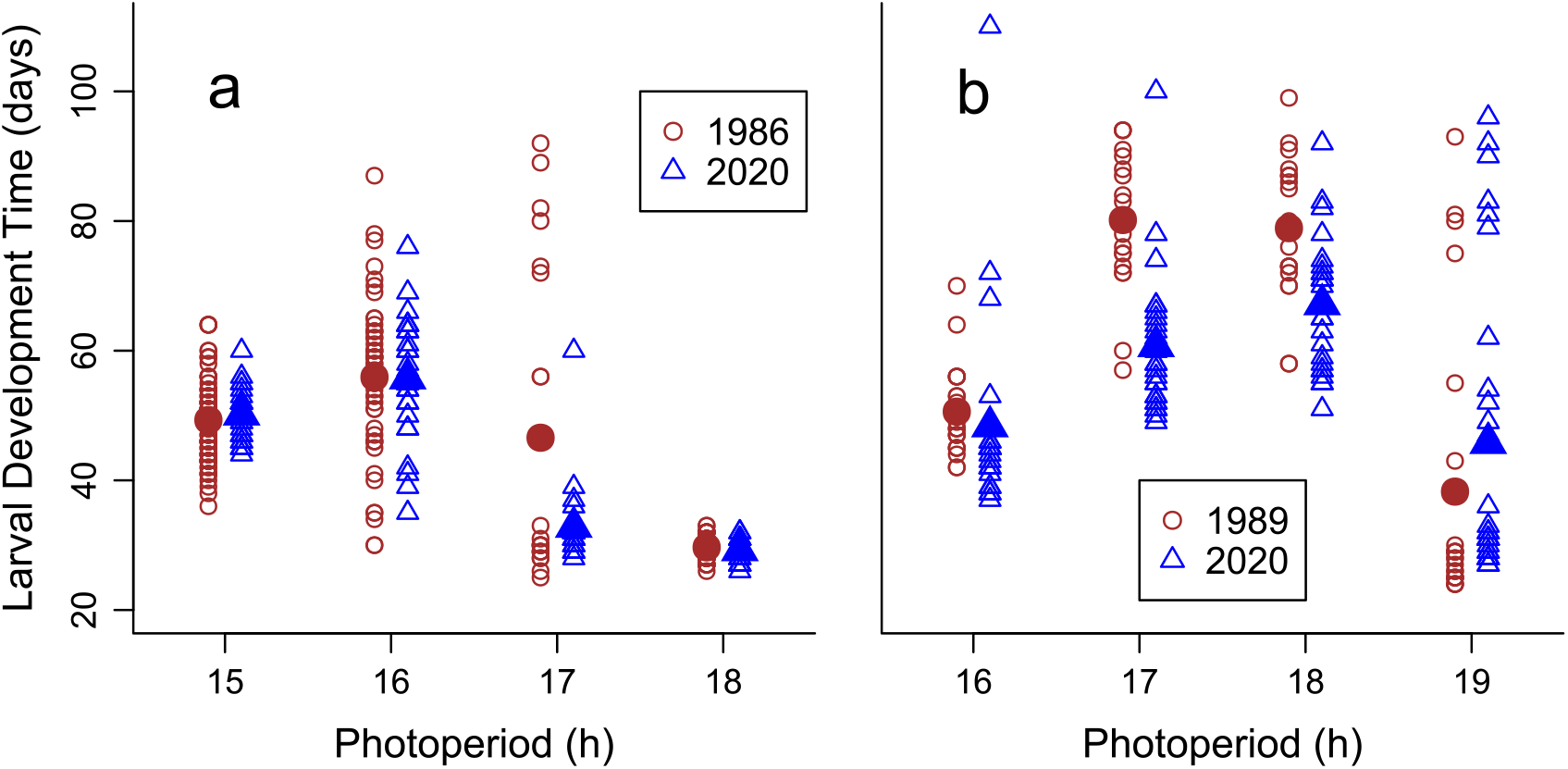
Comparison of larval development time of *P. aegeria* from historic (red circles) and contemporary (blue triangles) populations when raised at different photoperiods. (a) Bivoltine population from southern Sweden, historic population sampled from 1985-1987 (n=196), contemporary population sampled in 2020 (n=114). (b) Univoltine population from central Sweden, historic population sampled in 1989 (n=84), contemporary population sampled in 2020 (n=108). Small, open shapes represent individual data points. Large, closed shapes represent treatment means. Note differing photoperiod treatments used for each study site.

For the central population, we measured larval development time across four photoperiod treatments from 16L:9D to 19L:6D, comparing the results for 108 second-generation descendants of the contemporary population (2020) to 84 larvae measured in the historic experiments (1985-1989). In our final model, the response to photoperiod differed clearly between the historic and contemporary studies (Figure 3B, photoperiod x time-period: *F_3,183_*=7.20, *p*=0.00013). At 16L:8D, the shortest photoperiod, mean development time changed little between time-periods; however, it became substantially faster at the intermediate photoperiods historically associated with delayed larval development. At the longest photoperiod, which historically was the transition point between direct and delayed development, it instead became slower on average, associated with an increased number of individuals with long, delayed development. Only the difference at 17L:7D was statistically clear in a post-hoc test (Table S4), but it should be kept in mind that the multiple-comparison adjustment used here assumes tests are independent and is highly conservative. There was also a weak but unclear overall trend toward slower development in males (3.8 ± 2.2 days, *F_1,183_*=2.92, *p*=0.089).

We also used the same experiments to compare the incidence of pupal diapause between the historic and contemporary studies (southern 1985-87: *n*=192; southern 2020: *n*=114; central 1989: *n*=84; central 2020: *n*=107). For the southern population, the historic and contemporary reaction norms were highly similar (Figure 4A,B). There was no variation in developmental pathway at most photoperiods; all individuals chose the same pathway in both time periods. For 16L:8D, the one photoperiod with variation in diapause within time period, Fisher’s exact test did not find statistically clear evidence for a difference in diapause frequency between time periods (odds ratio=0.677, *p*=0.58). For the central Swedish population, frequency of diapause was also similar between the contemporary and historic studies at all photoperiods (Figure 4C,D). In this case only one of four photoperiods lacked variation in diapause (17L:7D: all diapause), but for the other three there was no statistically clear difference in diapause frequency between time periods by Fisher’s exact test (16L:8D: odds ratio=0, *p*=0.11; 18L:6D: odds ratio=0.64, *p*=1.0; 19L:5D: odds ratio=1.72, *p*=0.71).

**Figure 4.**
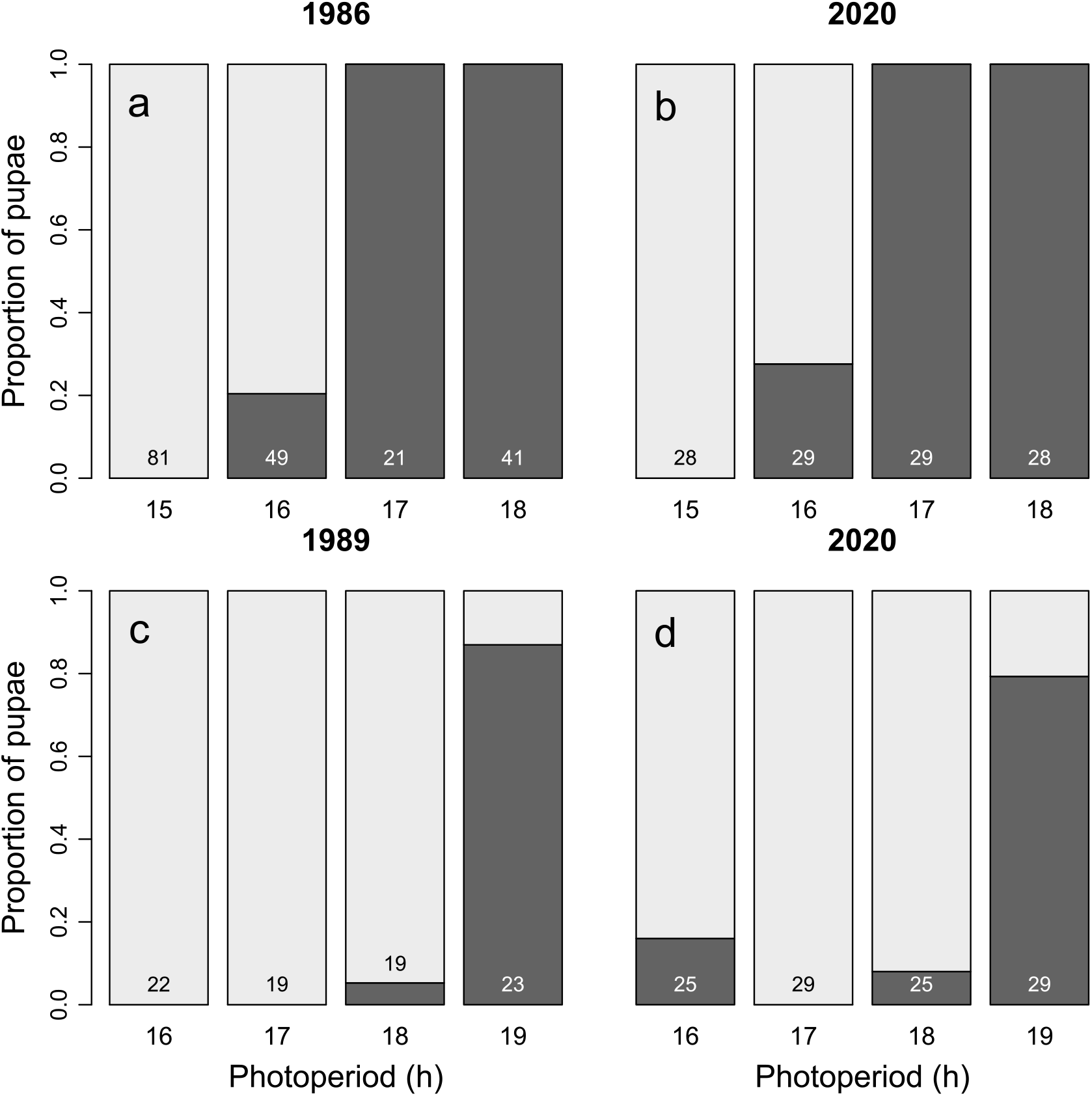
Comparison of frequency of pupal diapause versus direct development of *P. aegeria* from historic (A,C) and contemporary (B,D) populations when raised at different photoperiods. Light gray indicates diapause (>20 days as a pupa) and dark gray indicates direct development (<=20 days as pupa). Numbers at the bottom of each bar give sample size for that treatment and population combination. (A,B) Bivoltine population from southern Sweden. (C,D) Univoltine population from central Sweden. Note differing photoperiod treatments used for each study site.

Please see the supplemental materials for additional results on change in pupal mass between the historic and contemporary studies and for mixed model analysis of the effect of sex in the contemporary experiment.

## Discussion

By recreating some of the oldest experiments measuring the life history plasticity of multiple life stages, we found evidence for evolution of the photoperiodic reaction norms underlying the phenology of *P. aegeria* over the past 3 decades. This evolution, however, only occurred in the reaction norm for larval development time, not pupal diapause, despite substantial, well established spatial variation in both traits demonstrating their potential to evolve (19, 20, 24), Further, the evolution we did observe varied between populations, matching our prediction in the bivoltine population in southern Sweden, but taking a different form in the univoltine population in central Sweden.

As noted, the southern population displayed the predicted leftward shift in the reaction norm for larval development time, specifically increasing the frequency of rapid, direct development at the transition photoperiod for the historic population. This should allow this population to take advantage of the one month longer growing season they are now experiencing. This evolutionary change may allow them to add a third generation in some years, in line with the increased voltinism observed in a variety of Lepidopterans (28, 29). This change was stronger for males than females, but this may not reflect a difference in selection by climate change between the sexes. At the transition photoperiod, more males delayed their development than females in the historic study, a consequence of protandry in Swedish *P. aegeria* selecting for males to delay their development and enter diapause at longer photoperiods than females (30, 31). This difference between sexes may still exist in the contemporary population, but at a slightly shorter photoperiod not tested in these studies.

The change in the central Swedish population’s reaction norm was more complicated, but generally not aligned with our prediction. Development during the late-summer delay became faster (shifting this rate closer to that observed in the southern population), but at the transition photoperiod for the historic population, there is now a slightly higher frequency of individuals delaying their development, the opposite of the change in the southern population. Selection by climate change on phenology and associated plastic traits may be weaker for the central population. Despite similar mean warming, less of the observed climate change occurred during the growing season at the central site, which could reduce selection on the traits underlying phenology relative to the southern site. Nevertheless, spring is still occurring earlier in this population, and the delay in larval development time plays a key role in maintaining this population’s univoltine phenology despite variation in spring timing (20). The direct development pathway at long photoperiods has likely been under relaxed selection in the past (32), but the warmer and earlier springs they now experience could expose contemporary larvae to longer photoperiods than before. If warming does not give the central population enough time to complete a second generation, then selection should favor delayed larval development at longer photoperiods to maintain univoltinism. Regardless of the exact mechanism causing the difference, our results align with previous studies that have found variation in climate-associated evolution among populations (15, 33). Combined they demonstrate the key point that the evolutionary consequences of climate change often vary dramatically within a species’ range, so results from a single population cannot automatically be extrapolated to the entire species.

Although we found substantial evidence for evolutionary change in photoperiodic plasticity in larval development time, we found no evidence for analogous changes in pupal diapause. There was so little variation in this trait at most photoperiods that our preregistered statistical analyses could not be successfully applied to this data (Box 1), and further exploratory analyses also found no evidence for differences in this reaction norm. This lack of change could be due to the evolution of the larval reaction norm. The main proposed function of the late-summer larval developmental delay is to ensure that pupation occurs at an appropriate time to start diapause (20, 23). Thus, if the larvae adequately adjust this timing, it may mitigate the need for further selection on the pupal reaction norm. This result differs strikingly from the known spatial evolution of this trait in *P. aegeria,* in which latitudinal variation in the relationship between photoperiod and climate have led to well documented local adaptation of the reaction norm for pupal diapause, even over distances of less than 150 km (19, 20, 24). This extreme difference in results between spatial and temporal evolution suggests the popular space-for-time substitution (see 7, 8, 34) may not be appropriate for predicting evolutionary responses to climate change in all cases.

### Box 1. An example of preregistration in ecology and evolution

Preregistration originates in psychology and clinical trials as a method to distinguish *a priori* hypotheses generated before conducting research from later exploratory research after data has been collected, and it is advocated as a way to avoid under reporting of non-significant results and (often unconscious) manipulation of inferential statistics to artificially inflate p-values (39). In recent years, the preregistration discussion has begun in ecology and evolution (40, 41), but few empirical studies have been preregistered in these fields so far (42). Our study represents an early example of preregistration in the field and can provide guidance for the strengths and weaknesses of the approach in ecology and evolution.

Of course, the historic data used in our study existed and had been analyzed prior to preregistration. Instead, we preregistered our hypotheses and statistical methods before the collection of the new contemporary data (https://doi.org/10.17605/OSF.IO/6XQ4R). Preregistration makes clear which of our hypotheses were registered a priori and increases confidence in the associated statistics, but our study also illustrates a major limitation of preregistration for evolution and ecology. We used a standard experimental design and preregistered statistical methods that are often used to analyze the same traits in our study species (19, 20); nevertheless, our preregistered statistics for pupal diapause did not produce meaningful results. Ours is not the only study required to deviate from its registered analysis plan due to statistical issues (42), and other studies in ecology and evolution often involve much more complex data sets, study designs, and statistics than this study. Final data sets often have constraints that weren’t or couldn’t be anticipated at the time of preregistration, and thus deviating from preregistered data analysis plans may be the rule rather than the exception in ecology and evolution. While this challenge shouldn’t dissuade from using preregistration in ecology and evolution, particularly for hypotheses or study designs, we must also be mindful of the limits of preregistration and allow for the unforeseen complications that so often arise in these fields.

In addition to the evolution of the photoperiodic reaction norm for larval development time that we found, other evolutionary changes we have not yet examined could further adjust the phenology of *P. aegeria* in response to climate change. First, at photoperiods shorter than those included in our study, *P. aegeria* enters larval diapause instead of pupal diapause (23), and this transition could also be evolving in response to climate change. Second, while photoperiod is the main cue for seasonal plasticity in *P. aegeria,* it is modified by temperature, with higher temperatures shifting both larval and pupal reaction norms leftward toward direct development at shorter photoperiods (20). This temperature sensitivity could also have evolved, and a stronger plastic response to temperature may become beneficial if continued climate change decreases the reliability of photoperiod as a cue. Third, photoperiod is not constant in nature, and changing photoperiods can produce different results from constant photoperiods for the evolution of photoperiodic plasticity (e.g., 35). In the past, *P. aegeria* may not have needed this sensitivity because nearly all larval and pupal development occurred after the summer solstice; however, as phenology shifts earlier in the spring, being able to distinguish between increasing and decreasing photoperiods may become increasingly important (36).

Establishing whether and how evolution is contributing to the many phenological changes we have observed under climate change is essential to predicting whether and how observed phenological shifts will continue with future climate change (9). This is not, however, a simple task because the observed phenology of an organism is often determined by many different decisions made across its life cycle (2, 18). By building on experimental results from more than 30 years ago, we show here that the evolutionary consequences of climate change can be completely different for life history plasticity during two different life stages, both of which make key contributions to the species’ phenology. In our study, it appears that evolution of phenotypic plasticity at an early developmental stage was potentially sufficient to compensate for climate change, removing the need for evolution of later stages. Future studies may thus miss important evolutionary responses to climate change if they only focus on a single seasonal decision, rather than the whole range of plastic decisions contributing to phenology. Overall, we have shown that plastic responses to photoperiod are not fixed constraints, but that they can evolve and in doing so potentially allow evolution to at least partially correct for the cue-environment mismatches created by climate change. While studies of the mechanisms underlying phenotypic shifts are often phrased as evolution *or* plasticity, our results and others (15–17) indicate that it is more relevant to think in terms of evolution *of* plasticity.

## Materials and Methods

### Study sites and organisms

At Ransvik (56.27 N, 12.43 E), the southern Swedish site, *P. aegeria* is historically bivoltine, with the first, smaller flight peak during May-June and a second, larger flight peak during July-August (25). We collected 10 females for use in this study from Ransvik on 22 and 26 July 2020, during the second flight peak. At Riala (59.6212 N, 18.4963 E), the central Swedish site, *P. aegeria* is historically univoltine with a single flight peak during May-June (24). We collected 10 females for use in this study from Riala on 9 June 2020.

### Environmental temperature change

We downloaded from the Swedish Meteorological and Hydrological Institute daily mean temperature records for the decade prior to each study. No nearby weather stations spanned both the historic and contemporary time periods, so instead we selected the nearest pair of stations within 10 km of each other (Southern Sweden, historic: Kullen (Lat: 56.3012, Long: 12.4540); Southern Sweden, contemporary: Nyhamnsläge (Lat: 56.2451, Long: 12.5401); Central Sweden, historic: Arlanda (Lat: 59.6600, Long: 17.9208); Central Sweden, contemporary: Stockholm-Arlanda Flygplats (Lat: 59.6269, Long: 17.9545)). All stations except for Nyhamnsläge had complete coverage for the period in question. For Nyhamnsläge, we approximated the temperature of the missing 4.1% percent of days by using the days for which we had data to fit a regression predicting temperatures at Nyhamnsläge from temperatures at Stockholm-Arlanda Flygplats (*a*=3.29°C, *b*=0.802, *t_3499_*=177, *p*<1*10^-15^, *r^2^*=0.899). We then used this regression to estimate the temperatures of the missing days based on the corresponding temperatures at Stockholm-Arlanda Flygplats.

Using the daily mean temperature data, for each year we calculated the mean annual temperature, the mean temperature of each meteorological season (Dec-Feb, Mar-May, Jun-Aug, Sep-Nov), and the start, end, and length of the growing season. To estimate the length of the growing season, we estimated the start of the growing season as the first day of the year when the temperature was above 6°C and remained so for at least 7 days, and we used the first day after the summer equinox when temperature was below 6°C and remained so for at least 7 days as the end of the growing season (37). We used a threshold of 6°C because previous studies indicated this as the lower limit for development in *P. aegeria* (20). For all climate measures, we compared historic and contemporary values for each site using Welch’s t-test which allows for unequal variances.

### Historical data

This study used data from an experiment conducted in 1985-1987 for the Ransvik population (23), and 1989 for the Riala population (24). The information from original data sheets was digitized for this study. For Ransvik (23), we specifically used the data associated with Figure 1 supplemented by additional data from Figure 7 as appropriate. For Riala (24), we used the data associated with figure 3. After excluding one individual with unknown sex from analysis, we had historic data for 196 larvae and 192 pupae from Ransvik and 84 larvae and pupae from Riala.

### Recreation of historical experiments

Our experimental procedure was designed to replicate Nylin et al. (23, 24) as closely as possible, with additional details not provided by the manuscripts provided by the authors.

The historic experiments were primarily conducted on F1 offspring of wild females, but this was not entirely consistent, and some measurements were potentially conducted on F2 offspring. We conducted our study on the F2 offspring of the wild-caught females from each population. First, we raised the offspring of the wild-caught females for a full generation in the laboratory. Because the wild adults were collected at different times of year, this F1 generation was raised at different temperatures to synchronize them so we could test the F2 generations at about the same time (within one week of each other). Specifically, Riala F1 was raised at 14°C and Ransvik F1 at 23°C. Because we did not directly compare these populations, any potential parental effects should still be controlled for within population. Both F1 generations were raised at 22L:2D photoperiod to ensure direct development. Ten F1 larvae were raised per family, although individuals were not tracked. Instead, they were mixed in cages at a density of 25 per cage and fed using plants of *Dactylus glomerata.* We then conducted our study on the F2 offspring of this F1 generation. Specifically, for each population, we collected 12 offspring from 10 F1 females per population and divided them among the population’s 4 test photoperiods (see below) in a split-brood design. This resulted in an initial sample size of 30 individuals per treatment and 120 total for each population in the contemporary study. Caterpillars were randomly assigned to photoperiod treatments within 24 hours of hatching, with the constraint that an equal number of caterpillars from each female were assigned to each treatment.

Caterpillars were raised in growth chambers (3 Termaks KB84000-L and 2 KBP6345-L) at subset of photoperiods used in the historic study. Because of the historic differences in the reaction norms for the traits of interest (larval and pupal development time) between these populations (24), we used slightly different set of photoperiods for each population. For Ransvik, these photoperiod treatments were 15:9, 16:8, 17:9, and 18:6 hrs light:hrs dark. For Riala, they were 16:8, 17:9, 18:6, and 19:5 hrs light:hrs dark. Temperature was kept a constant 17°C, the same for all treatments and the same as the historic study. Caterpillars were kept individually on fresh *Poa sp.* grass as food. The diet was reported as *Poa annua* in the original experiment; however, *Poa* hybridizes and is extremely difficult to identify to species so the actual species used in the original experiment is uncertain. We used wild collected grass from the same region of Stockholm (Norra Djurgården) as used in the historic experiments *(pers. comm.)* so it should be as similar as possible to the historic diet. Grass was kept fresh during the experiment by placing its roots in fertilized water, and it was replaced as needed with fresh grass. It is unlikely to be relevant to our results, but the cups used for the caterpillars were also the same ones as used in the historic experiment.

Caterpillars were monitored to determine pupation date, and thus larval development time. Individuals that died before pupation were excluded from all analyses (southern: 6 larvae; central: 12 larvae). Two days after pupation (to allow time to harden), pupae were weighted and sexed, then returned to the same incubator in separate cups without grass. Pupae were monitored to determine adult eclosion date, and thus pupal development time. We considered pupae to have entered diapause if pupal development lasted longer than 20 days, and we specifically checked all pupae at this point to ensure that they remained alive. Pupal development time is bimodal, and this threshold encompasses all individuals identified as directly developing in the original study (23). It has also been used to effectively distinguish diapause from direct-developing individuals in recent work on *P. aegeria* (20). The one central Swedish pupae no longer alive at this time was excluded from analysis for pupal diapause.

### Statistics

Our hypotheses, experimental design, and statistical analysis plan were preregistered (https://doi.org/10.17605/OSF.IO/6XQ4R) before collection of any new data from our contemporary experiment. This historical data, of course, already existed and had been analyzed, although not by our proposed methods. Our analyses of environmental temperature change (see below) were exploratory and were not preregistered. All statistical analyses were conducted in R (v4.0.3). All statistical analyses were conducted separately for each population (southern or central) but following the same procedures.

We used linear models to test for a change in the response of larval development time to photoperiod between the historic and contemporary experiments. Larval development time was the response variable, and developmental photoperiod (as a factor), time period (historic versus contemporary), and sex were the predictor variables. We used backwards model selection (threshold of p>0.1) to remove non-significant interactions from the model. We then used the emmeans package (v1.6.1, (38)) to apply pairwise post-hoc tests with a Šidák correction to determine at which levels of photoperiod development time clearly differed between time periods or levels of the interaction between time period and sex (if retained in the model). These analyses were conducted as preregistered with the exception that post-hoc tests used the Šidák correction instead of Tukey’s HSD because it was deemed more appropriate for our analysis, but the Šidák correction is generally a conservative method.

We had preregistered a similar analysis for pupal diapause, using a generalized linear model with a binomial distribution to test for changes in the response to photoperiod between the contemporary and historic time periods. This method has been commonly used for assessing change in photoperiodic reaction norms for diapause (e.g., 19, 20); however, it could not be successfully applied to our final data. This failure was likely primarily caused by multiple photoperiods with no variation in diapause pathway (all individuals either entered diapause or developed directly). In the southern population, it failed to detect any evidence even for the graphically clear and obvious effect of photoperiod on diapause frequency, and there were similar problems for the central population, with a particularly poor fit for the contemporary data. In its place, we performed Fischer’s exact tests for just the photoperiod treatments in which both developmental pathways occurred during at least one experiment (contemporary or historic).

## Acknowledgments

We would like to thank Pauline Caillault and Olle Lindestad for their assistance in conducting the study. This research was funded by the Swedish Research Council (grant VR 2017-04500 to KG) and the Bolin Centre for Climate Research at Stockholm University (grant to KG).

## References

1. C. Parmesan, G. Yohe, A globally coherent fingerprint of climate change impacts across natural systems. Nature 421, 37–42 (2003).

2. S. J. Thackeray, et al., Phenological sensitivity to climate across taxa and trophic levels. Nature 535, 241–245 (2016).

3. Q. Ge, H. Wang, T. Rutishauser, J. Dai, Phenological response to climate change in China: a meta-analysis. Glob. Change Biol. 21, 265–274 (2015).

4. H. M. Kharouba, et al., Global shifts in the phenological synchrony of species interactions over recent decades. Proc. Natl. Acad. Sci. 115, 5211–5216 (2018).

5. S. S. Renner, C. M. Zohner, Climate Change and Phenological Mismatch in Trophic Interactions Among Plants, Insects, and Vertebrates. Annu. Rev. Ecol. Evol. Syst. 49, 165–182 (2018).

6. J. Merilä, Evolution in response to climate change: In pursuit of the missing evidence. BioEssays 34, 811–818 (2012).

7. J. Merilä, A. P. Hendry, Climate change, adaptation, and phenotypic plasticity: the problem and the evidence. Evol. Appl. 7, 1–14 (2014).

8. M. Kelly, Adaptation to climate change through genetic accommodation and assimilation of plastic phenotypes. Philos. Trans. R. Soc. B Biol. Sci. 374, 20180176 (2019).

9. H. E. Chmura, et al., The mechanisms of phenology: the patterns and processes of phenological shifts. Ecol. Monogr. 89, e01337 (2019).

10. W. E. Bradshaw, C. M. Holzapfel, Evolution of Animal Photoperiodism. Annu. Rev. Ecol. Evol. Syst. 3, 1–25 (2007).

11. R. A. Hut, S. Paolucci, R. Dor, C. P. Kyriacou, S. Daan, Latitudinal clines: an evolutionary view on biological rhythms. Proc. R. Soc. B Biol. Sci. 280, 20130433 (2013).

12. C. B. Edwards, L. H. Yang, Evolved Phenological Cueing Strategies Show Variable Responses to Climate Change. Am. Nat. 197, 16 (2021).

13. H. Van Dyck, D. Bonte, R. Puls, K. Gotthard, D. Maes, The lost generation hypothesis: could climate change drive ectotherms into a developmental trap? Oikos 124, 54–61 (2015).

14. N. Z. Kerr, et al., Developmental trap or demographic bonanza? Opposing consequences of earlier phenology in a changing climate for a multivoltine butterfly. Glob. Change Biol. 26, 2014–2027 (2020).

15. W. E. Bradshaw, C. M. Holzapfel, Genetic shift in photoperiodic response correlated with global warming. Proc. Natl. Acad. Sci. 98, 14509–14511 (2001).

16. T. Gomi, M. Nagasaka, T. Fukuda, H. Hagihara, Shifting of the life cycle and life-history traits of the fall webworm in relation to climate change. Entomol. Exp. Appl. 125, 179–184 (2007).

17. M. E. Nielsen, J. G. Kingsolver, Compensating for climate change–induced cue-environment mismatches: evidence for contemporary evolution of a photoperiodic reaction norm in Colias butterflies. Ecol. Lett. 23, 1129–1136 (2020).

18. B. Helm, B. M. Van Doren, D. Hoffmann, U. Hoffmann, Evolutionary Response to Climate Change in Migratory Pied Flycatchers. Curr. Biol. 29, 3714–3719.e4 (2019).

19. I. M. Aalberg Haugen, K. Gotthard, Diapause induction and relaxed selection on alternative developmental pathways in a butterfly. J. Anim. Ecol. 84, 464–472 (2015).

20. O. Lindestad, C. W. Wheat, S. Nylin, K. Gotthard, Local adaptation of photoperiodic plasticity maintains life cycle variation within latitudes in a butterfly. Ecology 100 (2019).

21. T. G. Shreeve, The effect of weather on the life cycle of the speckled wood butterfly Pararge aegeria. Ecol. Entomol. 11, 325–332 (1986).

22. C. Wiklund, A. Persson, P. O. Wickman, Larval aestivation and direct development as alternative strategies in the speckled wood butterfly, Pararge aegeria, in Sweden. Ecol. Entomol. 8, 233–238 (1983).

23. S. Nylin, P.-O. Wickman, C. Wiklund, Seasonal plasticity in growth and development of the speckled wood butterfly, Pararge aegeria (Satyrinae). Biol. J. Linn. Soc. 38, 155–171 (1989).

24. S. Nylin, P.-O. Wickman, C. Wiklund, Life-cycle regulation and life history plasticity in the speckled wood butterfly: are reaction norms predictable? Biol. J. Linn. Soc. 55, 143–157 (1995).

25. C. Wiklund, M. Friberg, Seasonal development and variation in abundance among four annual flight periods in a butterfly: a 20-year study of the speckled wood (Pararge aegeria). Biol. J. Linn. Soc. 102, 635–649 (2011).

26. J.-L. Tison, et al., Signature of post-glacial expansion and genetic structure at the northern range limit of the speckled wood butterfly: Speckled wood butterflies genetics. Biol. J. Linn. Soc. 113, 136–148 (2014).

27. O. Lindestad, S. Nylin, C. W. Wheat, K. Gotthard, Local adaptation of life cycles in a butterfly is associated with variation in several circadian clock genes. Mol. Ecol. 31, 1461–1475 (2022).

28. F. Altermatt, Climatic warming increases voltinism in European butterflies and moths. Proc. R. Soc. B Biol. Sci. 277, 1281–1287 (2010).

29. J. Pöyry, et al., Climate-induced increase of moth multivoltinism in boreal regions: Climate-induced increase in moth multivoltinism. Glob. Ecol. Biogeogr. 20, 289–298 (2011).

30. C. Wiklund, P.-O. Wickman, S. Nylin, A sex difference in the propensity to enter direct/diapause development: a result of selection for protandry. Evolution 46, 519–528 (1992).

31. S. Nylin, C. Wiklund, P.-O. Wickman, E. Garcia-Barros, Absence of Trade-Offs Between Sexual Size Dimorphism and Early Male Emergence in a Butterfly. Ecology 74, 1414–1427 (1993).

32. I. M. Aalberg Haugen, D. Berger, K. Gotthard, The evolution of alternative developmental pathways: footprints of selection on life-history traits in a butterfly: Alternative developmental pathways. J. Evol. Biol. 25, 1377–1388 (2012).

33. J. K. Higgins, H. J. MacLean, L. B. Buckley, J. G. Kingsolver, Geographic differences and microevolutionary changes in thermal sensitivity of butterfly larvae in response to climate. Funct. Ecol. 28, 982–989 (2014).

34. J. Verheyen, N. Tüzün, R. Stoks, Using natural laboratories to study evolution to global warming: contrasting altitudinal, latitudinal, and urbanization gradients. Curr. Opin. Insect Sci. 35, 10–19 (2019).

35. T. Harada, S. Nitta, K. Ito, Photoperiodic changes according to global warming in wing-form determination and diapause induction of a water strider, Aquarius paludum (Heteroptera: Gerridae). Appl. Entomol. Zool. 40, 461–466 (2005).

36. S. Nylin, Induction of diapause and seasonal morphs in butterflies and other insects: knowns, unknowns and the challenge of integration. Physiol. Entomol. 38, 96–104 (2013).

37. S. M. Kivelä, P. Välimäki, K. Gotthard, Evolution of alternative insect life histories in stochastic seasonal environments. Ecol. Evol. 6, 5596–5613 (2016).

38. R. V. Lenth, emmeans: Estimated Marginal Means, aka Least-Squares Means. R package version 1.6.1. (2021).

39. B. A. Nosek, C. R. Ebersole, A. C. DeHaven, D. T. Mellor, The preregistration revolution. Proc. Natl. Acad. Sci. 115, 2600–2606 (2018).

40. T. H. Parker, et al., Transparency in Ecology and Evolution: Real Problems, Real Solutions. Trends Ecol. Evol. 31, 711–719 (2016).

41. R. E. O’Dea, et al., Towards open, reliable, and transparent ecology and evolutionary biology. BMC Biol., 19:68 (2021).

42. L. G. O’Neill, T. H. Parker, S. C. Griffith, Nest size is predicted by female identity and the local environment in the blue tit (Cyanistes caeruleus), but is not related to the nest size of the genetic or foster mother. R. Soc. Open Sci. 5, 172036 (2018).

