## Supplemental materials for "Evolution of seasonal plasticity in response to climate change differs between life-stages of a butterfly"

Matthew E. Nielsen, Sören Nylin, Christer Wiklund, Karl Gotthard

**Table S1.** Comparison of seasonal temperatures and growing season duration during the 10 years prior to the studies on the southern Swedish population.

| Measurement | Historic mean (1976-1985) | Contemporary mean (2010-2019) | Change | *t* | *df* | *p* |
|---|---|---|---|---|---|---|
| Mean annual temperature (°C) | 7.53 | 9.00 | 1.47 | 4.01 | 15.52 | 0.0011 |
| Mean spring temperature (°C) | 5.63 | 7.61 | 1.98 | 5.11 | 16.05 | 0.00010 |
| Mean summer temperature (°C) | 15.73 | 17.07 | 1.34 | 3.87 | 15.46 | 0.0014 |
| Mean fall temperature (°C) | 9.08 | 9.86 | 0.78 | 2.37 | 16.03 | 0.031 |
| Mean winter temperature (°C) | -0.81 | 0.88 | 1.70 | 1.94 | 15.07 | 0.071 |
| Growing season start date (Julian date) | 118.1 | 97.7 | -20.4 | -4.84 | 14.76 | 0.00023 |
| Growing season end date (Julian date) | 319.5 | 330.5 | 11.0 | 1.82 | 17.64 | 0.086 |
| Length of growing season (days) | 201.4 | 232.8 | 31.4 | 4.58 | 17.29 | 0.00025 |

**Table S2.** Comparison of seasonal temperatures and growing season duration during the 10 years prior to the studies on the central Swedish population.

| Measurement | Historic mean (1979-1988) | Contemporary mean (2010-2019) | Change | *t* | *df* | *p* |
|---|---|---|---|---|---|---|
| Mean annual temperature (°C) | 5.57 | 7.11 | 1.54 | 3.82 | 17.66 | 0.0013 |
| Mean spring temperature (°C) | 4.36 | 6.13 | 1.76 | 5.01 | 17.65 | 0.000097 |
| Mean summer temperature (°C) | 15.80 | 16.66 | 0.86 | 2.20 | 17.61 | 0.041 |
| Mean fall temperature (°C) | 6.49 | 7.35 | 0.86 | 2.07 | 16.84 | 0.054 |
| Mean winter temperature (°C) | -3.47 | -0.62 | 2.86 | 2.65 | 17.68 | 0.016 |
| Growing season start date (Julian date) | 119.8 | 109.2 | -10.6 | -2.65 | 16.50 | 0.017 |
| Growing season end date (Julian date) | 299.3 | 298.2 | -1.1 | -0.18 | 17.97 | 0.86 |
| Length of growing season (days) | 179.5 | 189 | 9.5 | 1.22 | 16.77 | 0.24 |

**Table S3.** Post-hoc test for pairwise differences in larval development time at each photoperiod between sexes and between contemporary and historic populations from south Sweden using a Šidák correction.

| Photoperiod | Comparison | Difference estimate (days) | SE | df | *t* | *p* |
|---|---|---|---|---|---|---|
| 15 | Historic Female - Contemporary Female | 1.183 | 2.61 | 294 | 0.454 | 1.0000 |
| 15 | Historic Female - | 3.255 | 2.02 | 294 | 1.610 | 0.9365 |

| | | | | | | |
|---|---|---|---|---|---|---|
| | Historic Male | | | | | |
| 15 | Historic Female - Contemporary Male | 0.902 | 3.41 | 294 | 0.265 | 1.0000 |
| 15 | Contemporary Female - Historic Male | 2.073 | 2.45 | 294 | 0.845 | 1.0000 |
| 15 | Contemporary Female - Contemporary Male | -0.281 | 3.68 | 294 | -0.076 | 1.0000 |
| 15 | Historic Male - Contemporary Male | -2.353 | 3.29 | 294 | -0.714 | 1.0000 |
| 16 | Historic Female - Contemporary Female | -3.025 | 2.99 | 294 | -1.010 | 0.9999 |
| 16 | Historic Female - Historic Male | -6.945 | 2.60 | 294 | -2.672 | 0.1747 |
| 16 | Historic Female - Contemporary Male | -3.696 | 3.06 | 294 | -1.208 | 0.9980 |
| 16 | Contemporary Female - Historic Male | -3.920 | 2.97 | 294 | -1.319 | 0.9933 |
| 16 | Contemporary Female - Contemporary Male | -0.671 | 3.38 | 294 | -0.199 | 1.0000 |
| 16 | Historic Male- Contemporary Male | 3.249 | 3.04 | 294 | 1.070 | 0.9997 |
| 17 | Historic Female - Contemporary Female | 6.913 | 3.40 | 294 | 2.035 | 0.6491 |
| 17 | Historic Female - Historic Male | -13.212 | 4.09 | 294 | -3.232 | 0.0323 |
| 17 | Historic Female - Contemporary Male | 10.923 | 3.57 | 294 | 3.061 | 0.0562 |
| 17 | Contemporary Female - Historic Male | -20.125 | 3.94 | 294 | -5.109 | <.0001 |
| 17 | Contemporary Female - Contemporary Male | 4.010 | 3.40 | 294 | 1.181 | 0.9986 |
| 17 | Historic Male- Contemporary Male | 24.135 | 4.09 | 294 | 5.904 | <.0001 |
| 18 | Historic Female - Contemporary Female | 0.615 | 3.00 | 294 | 0.205 | 1.0000 |
| 18 | Historic Female - Historic Male | 2.259 | 2.81 | 294 | 0.804 | 1.0000 |
| 18 | Historic Female - Contemporary Male | 3.214 | 3.41 | 294 | 0.941 | 1.0000 |
| 18 | Contemporary Female - Historic Male | 1.644 | 2.94 | 294 | 0.560 | 1.0000 |
| 18 | Contemporary Female - Contemporary Male | 2.599 | 3.52 | 294 | 0.738 | 1.0000 |
| 18 | Historic Male- Contemporary Male | 0.955 | 3.36 | 294 | 0.284 | 1.0000 |

**Table S4.** Post-hoc test for pairwise differences in larval development time at each photoperiod between contemporary and historic populations from central Sweden using a Šidák correction.

| Photoperiod | Historic-contemporary difference (days) | SE | df | *t* | *p* |
|---|---|---|---|---|---|
| 16 | 2.06 | 4.41 | 183 | 0.468 | 0.9833 |
| 17 | 18.90 | 4.47 | 183 | 4.231 | 0.0001 |
| 18 | 10.83 | 4.56 | 183 | 2.375 | 0.0723 |
| 19 | -8.20 | 4.23 | 183 | -1.937 | 0.2000 |

**Pupal mass (pre-registered)**

Statistical analysis

To test for any changes in pupal mass between the historic and contemporary experiments, we used the approach as for larval development time: fitting separate linear models for each population. Pupal mass was the response variable with developmental photoperiod (as a factor), time period (historic versus contemporary), and sex as the predictor variables. We used backwards model selection (threshold of $p>0.1$) to remove non-significant interactions from the model and pairwise post-hoc tests with a Šidák correction to determine at which levels of photoperiod mass clearly differed between time periods, or how mass differed between photoperiod treatments in the absence of an interaction. We had to exclude 19 historic individuals from the southern Swedish population and 33 from the central Swedish population from this analysis because we did not have data on their mass. For the central Sweden population, this unfortunately included all historic individuals in the 19L:5D photoperiod treatment, so this treatment was excluded from analysis entirely.

Results

For the southern Swedish population, we compared the pupal mass of 112 offspring of P. aegeria collected in 2020 to 177 larvae descended from parents collected between 1985 and 1987, using 4 photoperiod treatments. We found a statistically clear interaction between photoperiod and time-period (photoperiod x time-period: $F_{3,281}$=5.14, $p$=0.0017, Figure S1A). In the post-hoc tests, this interaction was driven primarily by the 16L:8D treatment, at which contemporary pupae were 14 mg larger than in the past (Table S5). Males were 18.7 mg lighter in the historic population (s.e = 2.29, $t_{281}$=8.16, $p=1.1*10^{-14}$), but evidence for a slight increase in the effect of sex between time periods was not statistically clear ($b$=-6.16 mg, s.e.=3.70, $F_{1,281}$=2.78, $p$=0.097).

For the central Swedish population, we compared the pupal mass of 53 offspring of P. aegeria collected in 2020 to 79 individuals descended from parents collected in 1989, using 3 photoperiod treatments. We found no statistically clear evidence for a difference in the effect of photoperiod between time-periods (photoperiod x time-period: $F_{2,125}$=1.71, $p$=0.19; Figure S1B). Instead, we found an overall 20.0 mg increase in the mass of pupa from the contemporary population ($F_{1,127}$=74.5, $p=2.2*10^{-14}$). Photoperiod had a clear independent effect on mass ($F_{2,127}$=9.16, $p$=0.00019); based on the post-hoc tests, this difference was due to greater mass at 18L:6D than the other treatments (Table S6). Males were 22.5 mg lighter, independent of treatment and time period ($F_{1,127}$=97.9, $p<10^{-15}$), a difference similar to that observed in southern Sweden.

### Discussion

In the preregistration for this study, we hypothesized that unless an evolutionary change in larval development time was compensated for by a change in growth rate, we would see changes in pupal mass between the contemporary and historic experiments that corresponded to any observed changes in larval development time. As with larval development time, we found changes in body mass between the historic and contemporary experiments that differed greatly between populations. These changes did not, however, correspond to the changes in larval development time in the same population. In southern Sweden, pupal mass primarily differed between contemporary and historic experiments at a single photoperiod, but a different photoperiod from the one at which larval development time changed. In central Sweden, pupal mass increased independent of photoperiod, although we were not able to compare masses for the longest photoperiod treatment.

In some ways, it is not surprising that the changes and growth rate and development time do not correspond because the delay in larval development time at intermediate photoperiods decouples growth rate and development time (1). Development time is only likely to constrain body size growth rate is at its maximum limit, which is not the case during delayed development. Ultimately, it is difficult to be certain what has caused the observed changes in body size, but these changes appear to be independent of the changes in larval development time.

Though pupal mass is less affected by photoperiod than development pathway (1, 2, results of this study), it is likely to me more sensitive than development pathway to other factors, particularly diet quality. Because diet quality and other factors should be consistent within experiment (historic or contemporary), we can be more confident that changes in the response to photoperiod have heritable basis, but the overall increase in mass observed in central Sweden could instead be caused by a difference in diet quality between the historic and contemporary experiments and that result should be interpreted more conservatively.

**Mixed-effects model for contemporary data (pre-registered)**

#### Statistical analysis

To more accurately evaluate any differences in larval development time between males and females in the contemporary population, we fit a linear mixed model to just the contemporary data, accounting for the family-level structure in this data which was not available for the historic data. It should be noted that family structure is at the level of shared F1-parents and wild family structure was not tracked. Larval development time was the response variable, photoperiod (as a factor), sex, and their interaction were fixed effects, and family a random effect. We used backwards model selection with a threshold p-value of .1 calculated by Likelihood Ratio Test. To determine levels of photoperiod or its interaction with sex significantly differed, we applied post-hoc tests generating p-values using the Kenward-Roger method for degrees-of-freedom approximation and Tukey's correction for multiple tests. For pupal diapause, we attempted to perform the same analysis, except using a generalized linear mixed model with a binomial distribution and photoperiod as a continuous variable. Models were fit using lmer and glmer of the lme4 package (v1.1-26, (3)).

#### Results

For southern Sweden, we measured the larval development time of 114 offspring across 10 families from the contemporary (2020) population. Photoperiod treatment had a clear effect on development time in this population ($\chi^2_3$=192, $p$<$10^{-15}$), but we found no clear evidence for an effect of sex ($\chi^2_1$=1.86, $p$=0.17). In the post-hoc test, development time clearly differed between all levels of photoperiod ($p$<0.05). For central Sweden, we measured the larval development time of 108 offspring across 10 families from the contemporary (2020) population. Photoperiod treatment had a clear effect on development time in this population ($\chi^2_3$=35.5, $p$=9.3*$10^{-8}$), but we found no clear evidence for an effect of sex ($\chi^2_1$=2.43, $p$=0.12). In the post-hoc test, the photoperiod treatments fell into two groups with different larval development times: development time did not clearly differ between 16L:8D and 19L:5D ($t_{103}$=0.636, $p$=0.92) or 17L:7D and 18L:6D ($t_{103}$=-2.01, $p$=0.19), but did clearly differ in all other comparisons (p<0.05).

As with the main analysis, we could not effectively fit the preregistered mixed models to our data for diapause. For southern Sweden, no models converged. For central Sweden, all versions of the model were singular with no variation associated with family.

Discussion

These results correspond to our findings from the primary analysis for the contemporary population, including the same errors for pupal diapause. We generally found extremely weak to no evidence for an effect of sex on larval development time. These results may prove useful for any future study seeking to replicate this one and test for continued adaptation to climate change by allowing an analysis which accounts for family structure (at least when studying larval development time).

References

1. S. Nylin, P.-O. Wickman, C. Wiklund, Seasonal plasticity in growth and development of the speckled wood butterfly, Pararge aegeria (Satyrinae). *Biol. J. Linn. Soc.* **38**, 155–171 (1989).

2. O. Lindestad, I. M. Aalberg Haugen, K. Gotthard, Watching the days go by: Asymmetric regulation of caterpillar development by changes in photoperiod. *Ecol. Evol.* **11**, 5402–5412 (2021).

3. D. Bates, M. Mächler, B. Bolker, S. Walker, Fitting Linear Mixed-Effects Models Using lme4. *J. Stat. Softw.* **67** (2015).

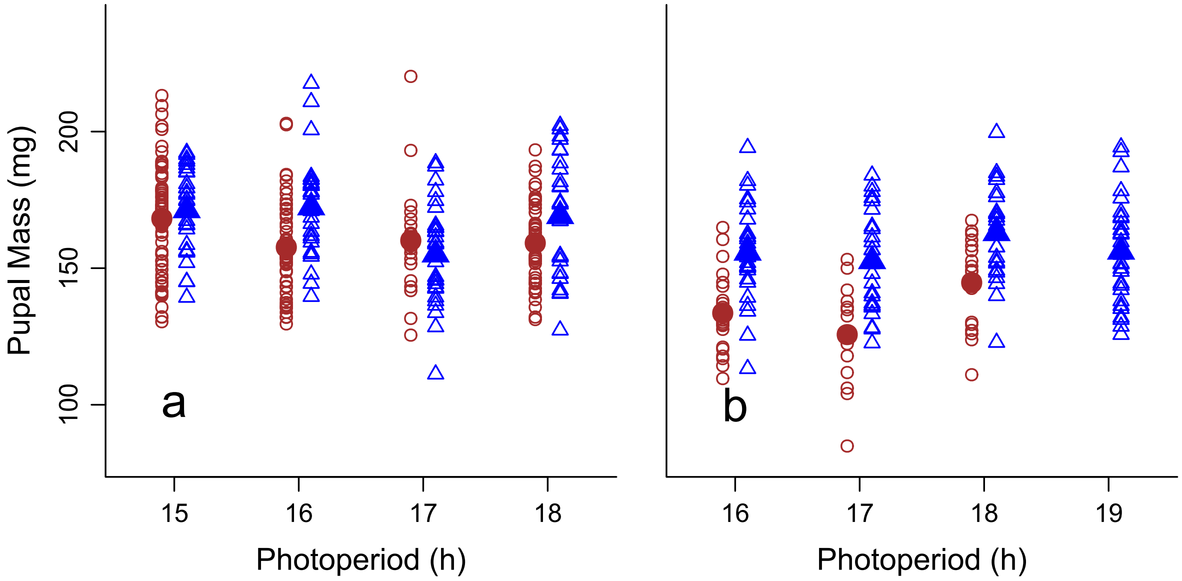


**Figure S1.** Comparison of pupal mass of *P. aegeria* from historic (red circles) and contemporary (blue triangles) populations when raised at different photoperiods. (a) Bivoltine population from southern Sweden, historic population sampled from 1985-1987, contemporary population sampled in 2020. (b) Univoltine population from central Sweden, historic population sampled in 1989, contemporary population sampled in 2020. No historic information on pupal mass was available from the 19L:5D photoperiod treatment for the historic experiment, so only contemporary data is plotted for this photoperiod, and it was excluded from analysis. Small, open shapes represent individual data points. Large, closed shapes represent treatment means.

**Table S5.** Post-hoc test for pairwise differences in pupal mass between contemporary and historic populations at each photoperiod from southern Sweden using a Šidák correction.

| Photoperiod | Historic-contemporary difference (mg) | SE | df | *t* | *p* |
|---|---|---|---|---|---|
| 15 | 2.66 | 3.45 | 281 | 0.771 | 0.9027 |
| 16 | -14.13 | 3.54 | 281 | -3.993 | 0.0003 |
| 17 | 3.45 | 4.40 | 281 | 0.783 | 0.8975 |
| 18 | -6.89 | 3.70 | 281 | -1.864 | 0.2306 |

**Table S6.** Post-hoc test for pairwise differences in pupal mass between photoperiod treatments for central Swedish population using a Šidák correction.

| Photoperiod comparison | Mass difference (mg) | SE | df | *t* | *p* |
|---|---|---|---|---|---|
| 16 - 17 | 2.41 | 2.76 | 127 | 0.875 | 0.7654 |
| 16 - 18 | -8.98 | 2.75 | 127 | -3.268 | 0.0042 |

| | | | | | |
|---|---|---|---|---|---|
| 17 - 18 | -11.39 | 2.82 | 127 | -4.044 | 0.0003 |
